# Towards model-based characterization of individual electrically stimulated nerve fibers

**DOI:** 10.1101/2025.07.21.665847

**Authors:** Rebecca C. Felsheim, David J. Sly, Stephen J. O’Leary, Mathias Dietz

## Abstract

Neuroprosthetics, such as cochlear implants or deep brain stimulators, can restore parts of the function of an impaired system. To improve such prosthetics, a detailed understanding of the electrical stimulation of nerve fibers is required. This knowledge can best be represented by computational models of the process. Currently, most models of individual electrically stimulated nerve fibers are based on many different datasets, which mainly consist of the average analysis values of recordings of many nerve fibers. While this is a valid approach for understanding the basic phenomena, both the combination of many different datasets and the average analysis can confound details in the response of the nerve fiber. To improve computational models of electrically stimulated nerve fibers further, we propose an optimization procedure that can fit the parameters of a neuron model to the response of a single nerve fiber to pulse-train stimulation. We show that in this way, the model can reproduce a wide variety of fiber responses of electrically stimulated auditory nerve fibers of guinea pigs in a remarkably detailed way on a scale of less than 1 ms. We analyze and discuss the certainty and generalizability of the parameter sets thus exposed. The model parameters found by the optimization procedure can then form the basis for a detailed fiber-by-fiber analysis, which we illustrate by a correlation analysis of the predicted phenomena (e.g., spike latency and refractory period) in the fiber response.

**Author Summary:** Neuroprosthetics can partially restore the function of an impaired neural system. Examples of such prosthetics are cochlear implants, which allow deaf people to hear, or deep-brain stimulators, which can reduce the tremor in Parkinson’s disease. While the mere existence of such prosthetics is already impressive, there is still room for improvement. Cochlear implant users, for example, have problems understanding speech in background noise, and deep-brain stimulators can have serious side effects, such as speech difficulties or limited fine motor control. To improve such implants, a detailed understanding of the underlying process, the electrical stimulation of nerve fibers, is required. A valuable representation of our understanding of the process is a computational model of electrically stimulated nerve fibers. Here, we optimized the parameters of one model, such that the behavior of 118 individual nerve fibers was represented, resulting in a single parameter set per fiber. In this way, a single model can reproduce a wide variety of response patterns, which allows for a detailed analysis of the individual fiber based on these parameters.

## Introduction

Neuroprosthetics, such as cochlear implants, stimulate neurons by generating an electric field that induces an electrical current. This can restore some of the functions of the impaired system, e.g., allowing deaf people to understand speech via cochlear implants [1] or reducing tremor in Parkinson‘s disease through deep-brain stimulation [2]. In many cases, however, the response characteristics differ from natural responses, leading to limitations, such as difficulties understanding speech in noise for cochlear implant users [1], or undesired side effects of deep-brain stimulators, such as speech difficulties or limited fine motor control [3].

A quantitative functional understanding of artificially stimulated neurons is of central importance for the safety, utility, and efficacy of neuroprosthetics. The most common way to better comprehend the behavior of such neurons is to perform single-fiber recordings in an experiment designed to answer a specific question. This data is then analyzed with respect to the question of interest (e.g., [4–9]).

To describe and combine knowledge about electrically stimulated neurons, computational models have been created, mostly based on the mean behavior of several single fiber recordings (e.g., [10–14]). Some models were based on a single set of recordings made for this purpose (e.g., [15]). However, most models, and especially the more recent and complex ones, were based on the analysis of many different single fiber recordings, collected for different purposes, and mostly published as the data from very few example fibers, or as the analysis of population mean values (e.g., [5,16]). Such analyses are valuable for understanding the principles of phenomena, such as the refractory period or facilitation, but the interdependence between different phenomena may be lost. It is also not appropriate to assume that the mean values represent the actual behavior of any “average” nerve fiber, as discussed by Marder and Taylor [17].

Collecting and analyzing data based on a specific research question has provided us with an invaluable understanding of the basic principles of spiking in electrically stimulated nerve fibers. These basic principles have been simulated or reproduced by many models (e.g., [10–12,18,19]). Some of these models already incorporated distributions of individual fiber parameters as they were reported in the task-specific experimental studies. However, the simulations of how these models respond to certain stimuli have never been experimentally validated. Neither has it been experimentally confirmed whether the model parameters obtained from different experimental studies result in a realistic model neuron, or even in a realistic distribution of neuron behaviors for a certain species. To achieve this, it is necessary to directly use the recordings from single nerve fibers to parameterize a model, resulting in one parameter set for each individual fiber. These parameters can then form the basis of a more detailed and diverse analysis of the behavior of the individual fiber. In addition, with these parameter sets, hypotheses about the results of new research questions can be created before conducting new experiments.

Existing work on fitting the behavior of each individual nerve fiber focused on fibers that were stimulated naturally via synapses, mostly via other nerve fibers (e.g., [20–25]). This comes with the disadvantage that the real-life input to the fibers is not entirely known. It is therefore approximated with white-noise stimulation, or a constant stimulus, in both cases often with a length of several seconds. In our work, the focus lies on fibers that are stimulated by neuroprosthetics, for which this approach has not previously been applied. Focusing on neuroprosthetics comes with the advantage that the input to the fibers is known and can be reproduced. Usually, these devices use pulse trains with a constant pulse shape, which are amplitude-or frequency-modulated to transmit information [2,26]. The pulsatile stimulation also allows the analysis of the relationship between each pulse and its immediate neural response, similar to the advantage of knowing the impulse response in systems theory.

The basis for the fitting procedure is formed by a model, and, as in many of the abovementioned studies [21–25], we use a generalized leaky-integrate-and-fire model. More specifically, we use the adaptive Leaky-Integrate and Firing Probability model (aLIFP, [27]), which not only models a wide range of spike interaction phenomena, but also includes a detailed simulation of the spike timing. The simulation of the spike timing includes both the latency between the stimulation and the spike, and the stochasticity of this latency, called jitter.

Fitting a stochastic model to stochastic data can be very tedious, as the metric quantifying the difference between the model must consider the stochasticity of the fiber responses. To avoid this problem, we use data in which all stimuli have been repeated multiple times, which allows us to obtain information about the amount of stochasticity of the nerve fiber. Also, the aLIFP model does not describe the stochasticity of the process with a random output. Instead, the output is the spike probability as a function of time, which describes the stochasticity with a deterministic output. In this way we can compare the recordings of the nerve fibers and the model output and gain information on the amount of stochasticity. The combination of the deterministic output of the aLIFP model and the detailed description of the fiber response allows us to fit the responses of each nerve fiber in detail.

The aim of this work was to separately fit the parameters of the aLIFP model [27] to each neuron in a set of in-vivo recordings of electrically stimulated auditory-nerve fibers in guinea pigs, as recorded by Heffer et al. [9]. Each fiber was stimulated with different pulse rates and amplitudes, each condition being repeated several times. After fitting the model to each fiber of the in-vivo recordings, we analyzed the quality and consistency of the fit. Finally, we demonstrate the advantages of this method by evaluating relationships between the different phenomena in the behavior of the nerve fiber.

## Results

The data used in this work was originally recorded by Heffer et al. [9]. They recorded the responses of guinea pig auditory nerve fibers to constant amplitude pulse trains of 200 pps, 1000 pps, 2000 pps, and, for some fibers, 5000 pps. Each pulse train was presented at multiple amplitudes, and each condition was repeated multiple times. A more detailed description of the data can be found in the Methods section and in the original publication [9].

The model used in our optimization procedure was the adaptive Leaky-Integrate and Firing Probability (aLIFP) model [27], which is based on the sequential biphasic leaky integrate and fire model [10]. The phenomenological aLIFP model predicts the spike times in response to the stimulation with electrical pulses. However, instead of predicting exact and stochastic time points, the model response is the spike probability over time. In this way, the stochasticity of the nerve response can be described with a deterministic model output. This approach forms the basis for this work, as the fitting of the model parameters is facilitated through the deterministic response while still being able to describe the details, including the spike time variance. The aLIFP model includes spike latency and jitter, as well as the pulse interaction phenomena refractoriness, facilitation, accommodation, and adaptation. A more detailed description of these phenomena can be found in the Methods section and in the original publication of the model [27].

During the optimization procedure, the model parameters were adjusted such that the spike rate of the model in response to stimulation with 200 pps, 1000 pps, and 2000 pps is as similar to the data as possible. To quantify the difference of the model response compared to the data, three similarity terms were used. The first one computed the normalized cross-correlation (xcorr) between the first part of the response of the model and the data. A value of 1 indicated a perfect correlation, and a value of 0 indicated no correlation at all. The second term was the complex vector strength difference (cVS_diff_), which quantified not only the difference in phase locking between the model and the data but also the difference in the phase position. A value of 0 indicated that the complex vector strength was the same in the model as in the data, which required the same degree of phase locking and the same phase position. The larger the difference between the data and model phase locking and position, the higher the value, up to a maximum of 2. The last similarity term was the absolute difference between the square root of the simulated and experimentally obtained rate in the second part of the signal (rate_diff_). Here, a value of 0 indicated complete equality, with rising values indicating a higher inequality. All three similarity terms were described in greater detail in the Methods section.

### Individual response patterns

When looking at the individual responses of the nerve fibers, a wide variety of response patterns to the same stimuli was seen. Figure 1 shows the responses of four exemplary nerve fibers to 200 pps, 1000 pps, and 2000 pps stimulation at two amplitudes each. These four fibers were chosen to show a variety of different response patterns and success in fitting the model response to the data. The nerve fiber responses are shown in blue and the model simulations based on individually optimized parameters are shown in red.

**Figure 1:**
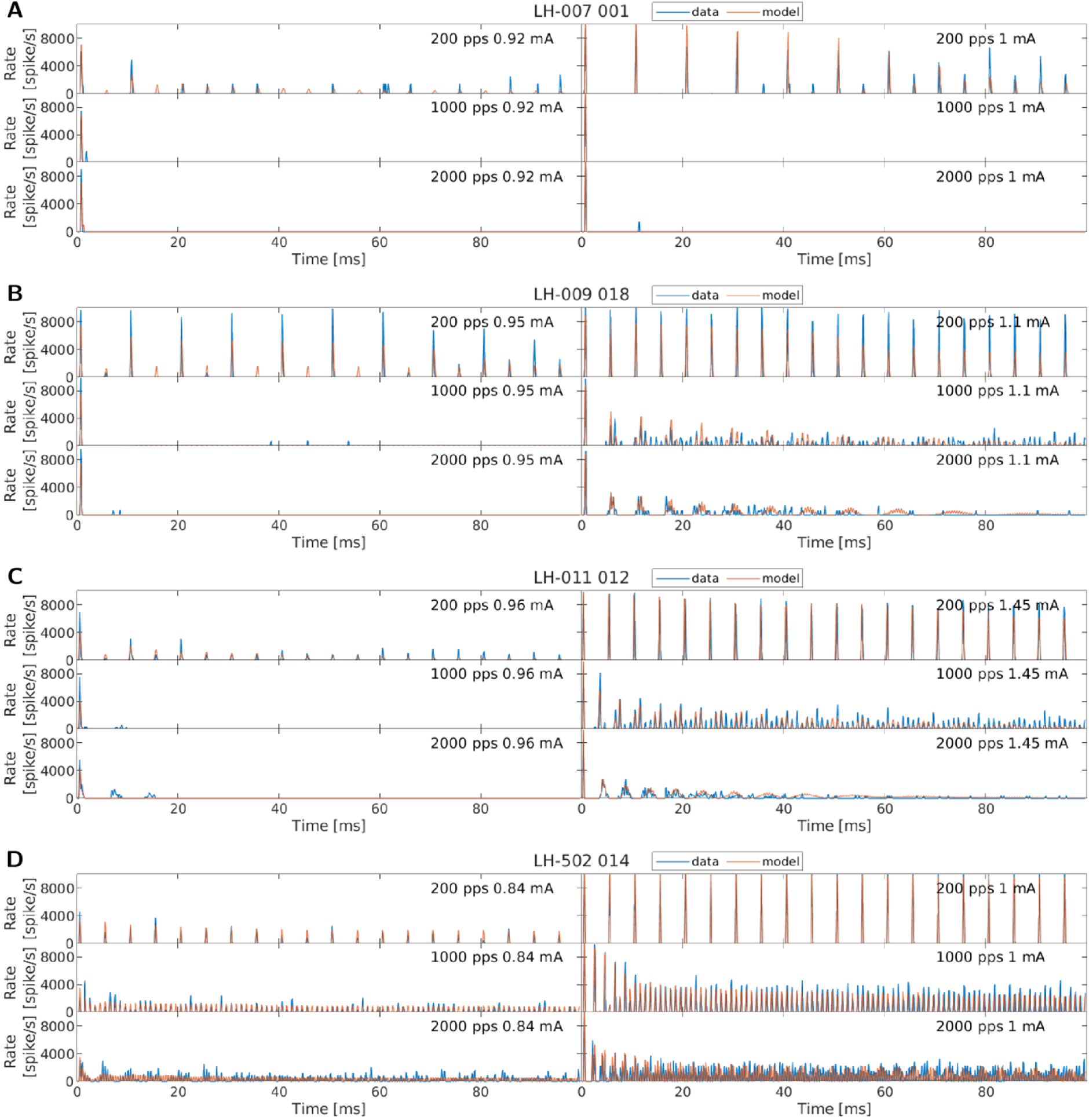
Recorded spike responses of four exemplary nerve fibers stimulated with 200 pps, 1000 pps, or 2000 pps, at two different amplitudes (blue), together with the model simulations based on the fitted parameters for the respective fiber (red).

Comparing the responses of the fibers to 200 pps stimulation, some fibers responded to every pulse (e.g., LH-011_012 in panel C, LH-502_014 in panel D), while others responded only to every second pulse (e.g., LH-007_001 in panel A). Another group of fibers responded to every second pulse at low amplitudes and to every pulse if stimulated with a higher current (e.g., LH-009_018 in panel B).

The response to higher pulse rates (1000 pps or 2000 pps) also varied greatly across fibers. The response to the first pulse was very similar across all fibers. Here, only the dynamic range, i.e., the stimulation amplitude range across which the response probability changes, was different for the individual nerve fibers. However, afterwards the patterns were quite different. Some nerve fibers did not respond or responded with a very low spike rate to the stimulation after the first pulse (e.g., LH-007_001 in panel A). Other nerve fibers exhibited a temporally modulated pattern with areas of a high response rate alternating with areas with a low response rate (e.g., LH-009_018 in panel B, LH-011_012 in panel C). This then slowly changed into a more random or less modulated response, with the transition again being fiber specific. Yet another group of nerve fibers exhibited a very high, less modulated response rate from the very beginning of the stimulation (e.g., LH-502_014 in panel D). The three patterns presented should not be seen as discrete groups into which the fibers can be classified, but as a continuous response space. Many nerve fibers exhibited responses that can best be described by a mixture of these patterns.

When looking at the data in Figure 1 and the model simulations in red, it can be seen that we were able to adjust the model parameters such that these patterns can be simulated with high accuracy and a high level of detail. This is also illustrated by the magnified snippets in Figure 2. However, when comparing the measured data with the model response closely, some limits of the model simulations were found.

**Figure 2:**
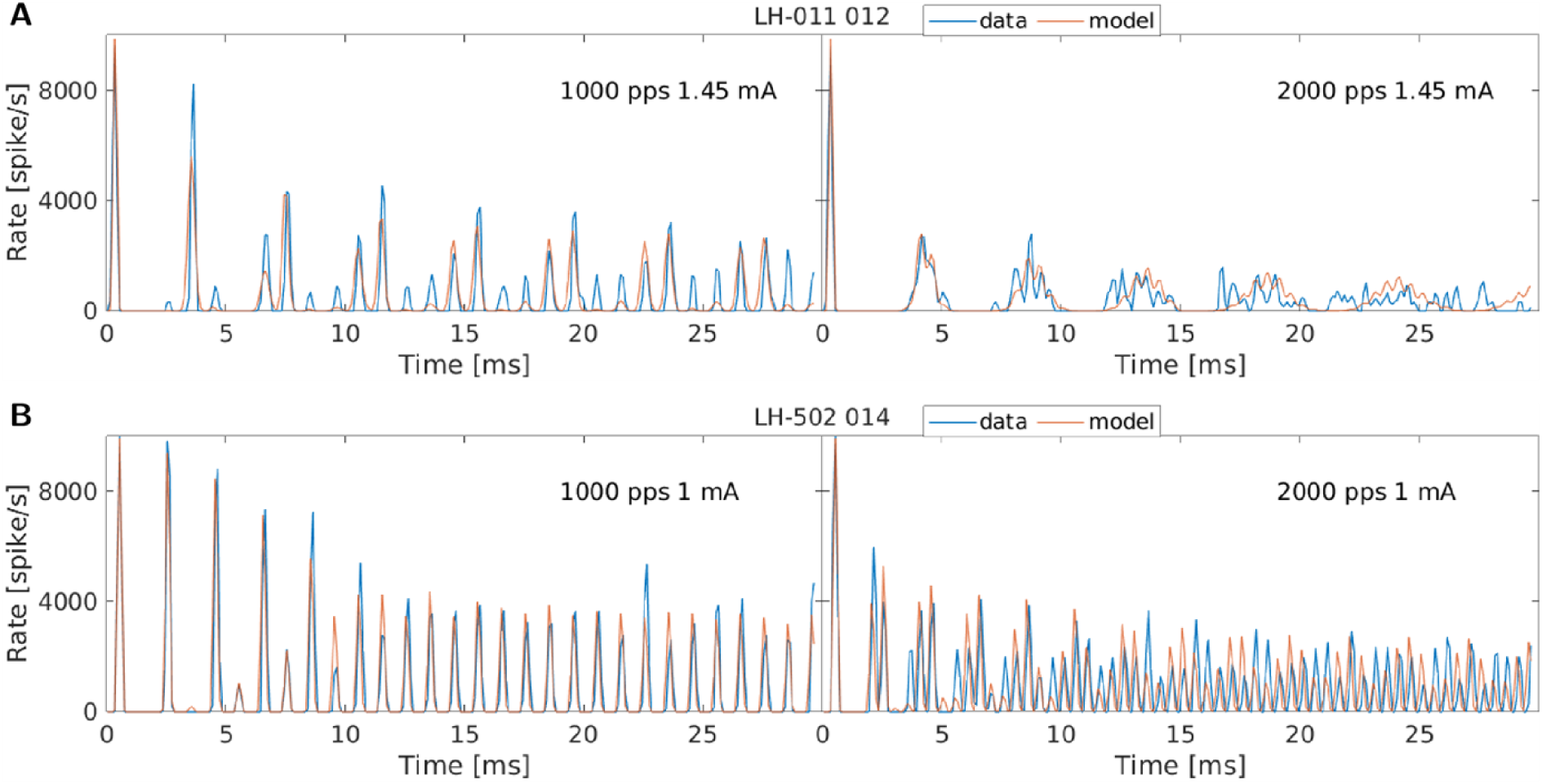
To illustrate the detail with which the aLIFP model can replicate the fiber responses, this figure shows the response to the first 30 ms of stimulation at two different pulse rates for the fibers LH-009_043 and LH-510_018 (Figure 1C+D).

The first limit that was observed is that the transition between a modulated response and a more stochastic or less modulated response was partially difficult to reproduce with the model. This can be seen in Figure 1 for fiber LH-009_018 stimulated with 1000 pps (panel B) or for fiber LH_011_012 (panel C) stimulated with 1000 pps. In general, the model tends to overestimate the modulation in the response.

Some fibers (e.g., LH-011_012 in panel C at 2000 pps) had a relatively high and modulated response for lower amplitudes at the beginning of the stimulation. This behavior could not be simulated with the used aLIFP model. However, more stochastic responses to stimulation with lower amplitudes (e.g., LH-502_014 in panel D at 1000 pps) could be modelled.

When looking at the responses at low response rates, it might look like the simulation of the model was less accurate compared to the responses at a higher rate (e.g., for fiber LH-502_014 in panel D at 2000 pps). However, a reduced spike probability was represented differently in the model response compared to the data. For the data, the stochasticity at a low response rate resulted in a spike every now and then. The response rate that was obtained from a single spike was limited by the number of repetitions of the stimulus. The model, on the other hand, was not limited to discrete spikes. Here the spike probability, and thus also the response rate, was an arbitrarily low number. A reduced spike probability was then described by a very low response rate over a longer interval. In such a case, the fiber responses and the model response might differ visually a lot but represent the same long-term response probability. This is also the reason why the cross-correlation was only computed at the beginning of the signal and why we focused on the average rate at the end.

### Overall fitting success

After discussing the individual response patterns of the different nerve fibers and the success in predicting them, we have evaluated the overall success of our parameter optimization. For this, we plotted the mean values of our three optimization metrics across all amplitudes but separately for each pulse rate in Figure 3. The values for 5000 pps in the rightmost boxplot are separated by a line, as this data has not been used during the fit but only for simulation purposes. The 5000 pps data and model simulations are also only based on the 64 fibers for which it was available, whereas the data from the other pulse rates is based on all 118 fibers. The values for the four fibers discussed above are marked with symbols.

**Figure 3:**
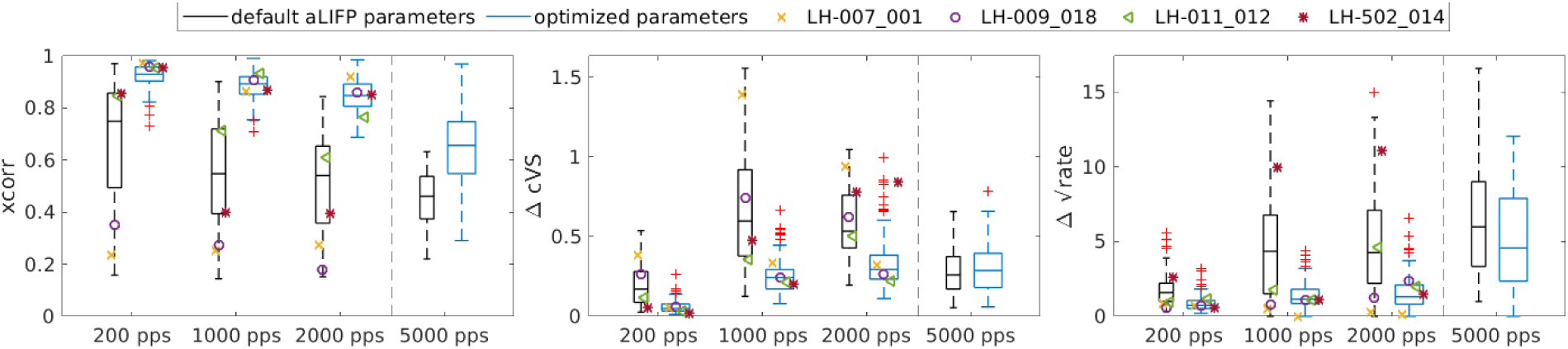
A comparison of the simulations with the optimized parameters in blue and the simulations using the default parameters of the aLIFP model in black to quantify the fitting success. Each panel shows one of the terms expressing the similarity between the model and the data: (A) cross-correlation (xcorr), (B) complex vector strength difference (ΔcVS), and (C) rate difference (Δ√rate), separately for each pulse rate, averaged across all amplitudes. The rightmost boxplot in each of the three panels gives the success of the simulation for the 5000 pps condition for all fibers where the data was available. However, this condition was not used during the optimization procedure. The values for the four exemplary fibers discussed above are marked, if available.

To provide a baseline for comparison with our fit metrics, we also simulated the responses of all nerve fibers with the default parameters of the aLIFP model. Only the membrane resistance and the threshold standard deviation were individualized to ensure a response over the range of stimulating amplitudes. These baseline simulations are shown in black in Figure 3, while the simulations with the individually fitted parameters are shown in blue. Comparing these two showed that while there are some fibers whose behavior can be predicted with the default parameters, fitting the parameters to each individual fiber drastically improved the simulations for 200 pps, 1000 pps, and 2000 pps. The cross-correlation and the rate difference also improved for 5000 pps when comparing the individualized parameters with the default parameters, but less than the fitted pulse rates. For the 5000 pps case, the ΔcVS did not change compared to the default values. This is likely due to the overall low vector strength at 5000 pps. The difference in the phase position then has a lower impact on the difference. Overall, it can be said that we successfully fitted the 200 pps, 1000 pps, and 2000 pps data. We have also measured the duration of the fitting procedure, which was 8 ± 4 hours per fiber on a single core of a Xeon Gold 6238 CPU.

Comparing the similarity terms across different pulse rates revealed that the similarity was reduced for the pulse rates higher than 200 pps, which is likely related to the increased complexity of the response. The cross correlation and the rate difference for the 5000 pps conditions values were worse than all other pulse rates, which indicates that the fitted values did not generalize to this much higher pulse rate. This can be explained by the fact that for the 5000 pps condition points on the pulse-interaction curves become relevant, for which no information was available previously. The difference in complex vector strength did not increase, which is due to an overall lower vector strength. It is also interesting to note that the worsening in the similarity term for 5000 pps did not happen for the default aLIFP model parameters, which indicates that the model might not be worse in simulating the 5000 pps data per se.

### Parameter certainty

To evaluate the precision of our parameter optimization procedure, we repeated the procedure 20 times for two fibers. The optimization consisted of five steps, which are described in detail in the Methods. In the third of these five steps the parameters describing the refractory period, facilitation, and accommodation were fitted. As our optimization problem contained many local optima, this optimization step was always started from six starting points, leading to six candidate results for the third optimization step. The remaining optimization steps were then based on the best candidate solution of the third step. In the default optimization the starting points for the third optimization step were manually selected and the same for every optimization. For the evaluation of the parameter certainty, we forced our algorithm to consider also other areas on the optimization landscape. To do this while avoiding entirely unrealistic solutions, we added a random vector to each of the six starting points. This random vector was drawn anew for every optimization started. The rest of the optimization remained the same.

To evaluate the optimizations, we computed characteristic values for the latency, jitter, duration of the refractory period, facilitation duration, accommodation duration, and adaptation amplitude. The latency is the mean duration between the pulse and the spike, and its standard deviation is the jitter. Both were characterized by their value at a spike probability of 0.5 in response to a single pulse. The refractory period is the duration of decreased excitability of the cell or an increase in spike threshold. It was described by the duration until the spike threshold reaches 1.5 times the resting threshold. If a pulse does not evoke a spike, the excitability is first increased, or the threshold decreased, which is called facilitation. Then, the excitability is reduced, or the spike threshold increased, which is called accommodation [8]. We characterized the facilitation by the duration of the threshold decrease and the accommodation by the time between the start of the threshold increase until the threshold decreased to again 1.01 times the resting threshold. Finally, the adaptation is characterized by the relative increase of the threshold 10 ms after a single pulse. As the optimization for the parameters describing the dynamic range of the fiber (membrane resistance and threshold standard deviation) were found using a deterministic procedure, these parameters and the dynamic range were the same for each run and were therefore not included in this analysis.

Fiber LH-007_001 was chosen as the first fiber to be refitted. As can be seen in Figure 1, the fiber exhibits close to no response after the first spike if stimulated with 1000 pps or 2000 pps. This causes a high degree of freedom during the optimization procedure, as it is conceivable that many parameter combinations result in a similar simulation. We chose this fiber on purpose to see how the optimization procedure would use this freedom. The result of this first refitting procedure is shown in Figure 4. Panels A-C show the values of the three similarity terms separately for each pulse rate. The green crosses mark the values of the original fit. It can be seen that the values across all repetitions were very similar, with the highest deviation in the rate difference. This shows that our procedure resulted in parameters that describe the recorded data similarly well, even if the starting points differed, which was expected due to this simple response pattern. Looking at the parameters shows differences larger than a factor of 2 (6 dB) across the 20 simulations (e.g., f_1_ and all refractory parameters), with the most extreme factor of 10^13^ (260 dB) relative to the original value for the adaptation time constant τ_a_ in a single run. The latency and jitter parameters, f_3_ of facilitation and accommodation, and c_a_ of adaptation were similar across all fits. Despite the high differences in parameter values, the characteristic values of the phenomena differ at most by a factor of 2 (6 dB). This indicates that the optimization procedure seems to explore the parameter space, but even the sparse spike pattern enforces strong enough constraints to get similar phenomena values. However, the high differences in the parameter values indicate a strong interaction between some model parameters.

**Figure 4:**
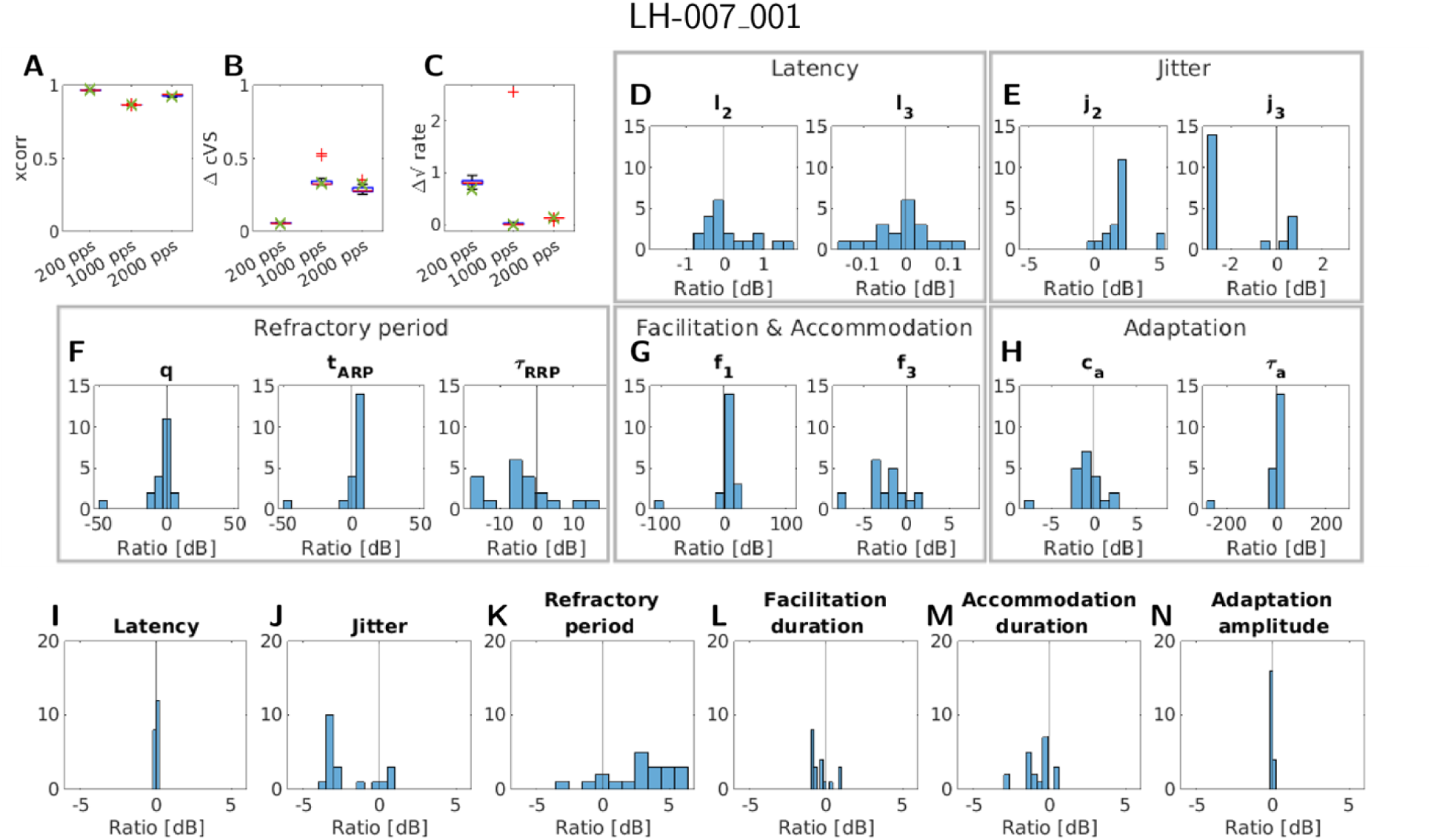
An overview of the results of the refitting procedure for the fiber LH-007_001. Panels A-C give the three similarity terms separately for each of the pulse rates considered. The green crosses mark the value of the original fit. In panels D-H the change in decibels relative to the original fit for each fitted parameter is shown, except for the membrane resistance and threshold standard deviation. Panels I-N give the change in decibels relative to the original fit for the characteristic values of each phenomenon, except for the dynamic range.

The second fiber (LH-009_018) for this evaluation was chosen to have a more complex response pattern. Nevertheless, the overall picture was similar, as can be seen in Figure 5. Comparing the similarity terms in the panels A-C of Figure 5 showed that the cross-correlation and the complex vector strength difference stay very similar to the original fit. However, the rate difference had a higher variability across the fits, especially at 2000 pps. This can be explained by the fact that the differences in the individual parameters or phenomena cumulate over stimulus duration, leading to higher differences at the end compared to the beginning. This then leads to a higher variability for the rate difference, which is computed at the end of the signal compared to the correlation computed for the first part. This was observed for the previously considered fiber due to its response close to zero for the higher pulse rates.

**Figure 5:**
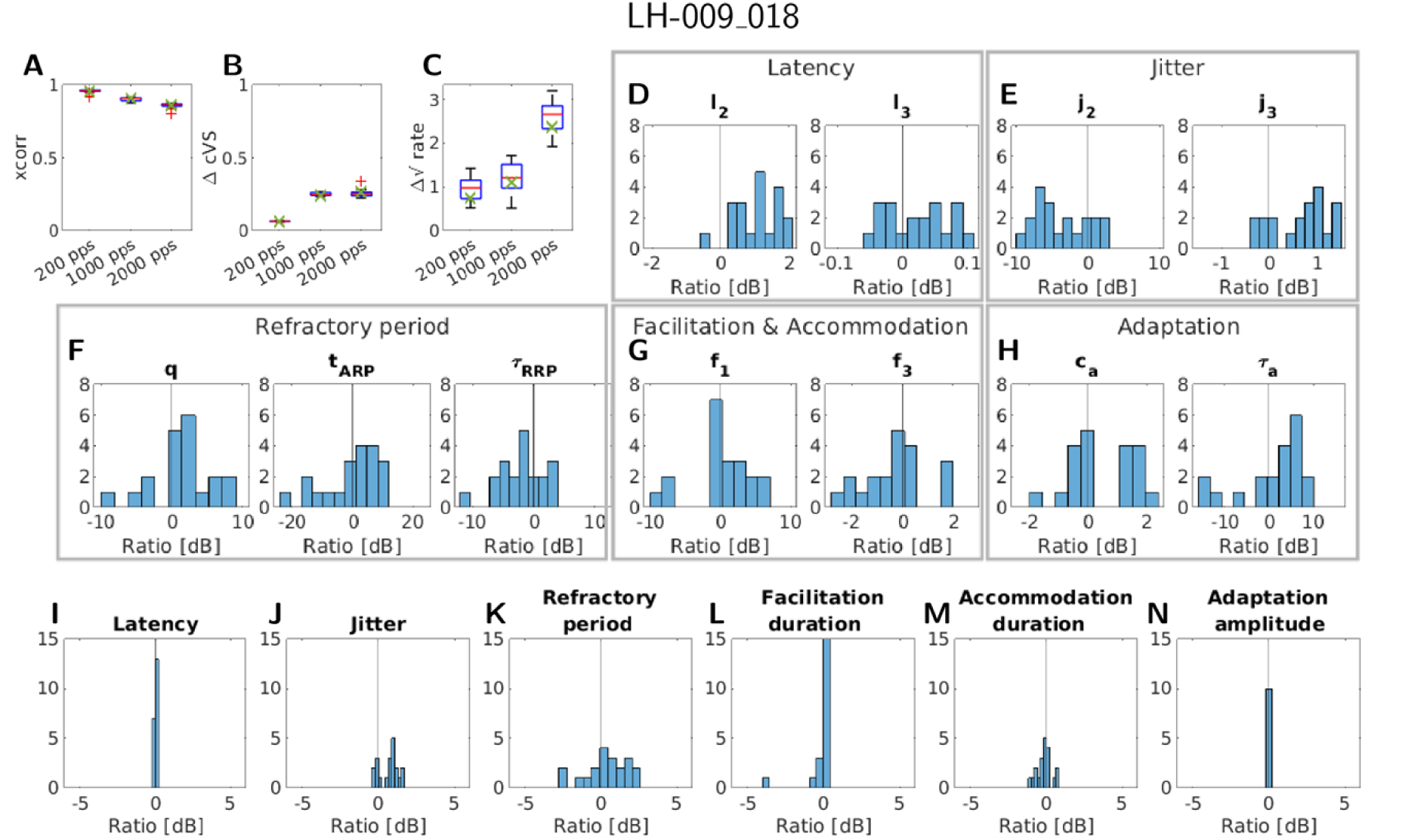
An overview of the results of the refitting procedure for the fiber LH-009_018. The panels A-C give the three similarity terms separately for each of the considered pulse rates. The green crosses mark the value of the original fit. Panel D-H shows the change relative to the original fit for each fitted parameter, except for the membrane resistance and threshold standard deviation. Panel I-N gives the change relative to the original fit for the characteristic values of each phenomenon, except for the dynamic range.

When looking at the parameter changes in panels D-H of Figure 5, it can be seen that the change in parameters was less extreme compared to fiber LH-007_001. This shows that the higher fiber response contains more information and therefore led to less ambiguity in possible parameter sets. A similar picture can be seen when looking at the change in characteristic value of the phenomena (Figure 5I-N). Here especially the range of jitter at 50% spike probability was much lower compared to fiber LH-007_001. Of all phenomena evaluated, the latency, jitter, and adaptation seemed to be the most stable.

Comparing the variability in the parameters to the variability in the phenomena for both nerve fibers showed that in both cases the phenomena exhibit less variability. This indicated, as mentioned above, that multiple parameter sets of the model led to similar response properties, even at the basis of individual phenomena. It might therefore be reasonable to use this knowledge to improve the number and type of (free) parameters in the aLIFP model.

### Stimulation with 5000 pps

Figure 3 shows that the parameters fitted to 200 pps, 1000 pps, and 2000 pps did not extrapolate well to simulate the 5000 pps data. This led to the question if the difficulties in predicting the 5000 pps data were due to a model limitation or due to the fitting procedure. To answer this question, we selected 10 fibers for which the 5000 pps data were also available and used all four pulse rates during the optimization procedure.

In the upper row of the two panels of Figure 6, the data and the model responses are shown for 1000 pps, 2000 pps, and 5000 pps at the highest stimulating amplitudes for the originally fitted parameters. This illustrates that the 5000 pps data could not be predicted by the original fit, while the responses to the trained pulse rates could be simulated. In the lower row of the two panels of Figure 6, the same data is shown, but the model simulations were based on the parameter sets that had also been fitted to the 5000 pps data. These fits show that the model was very well able to simulate the 5000 pps data if the data was included in the optimization procedure. They also show that the simulations of the 1000 pps and 2000 pps data did not get worse by including the 5000 pps data. For the fiber LH-022_004 in panel A, they even became slightly better. This might happen because the 5000 pps data helped to disambiguate between multiple possible parameter sets.

**Figure 6:**
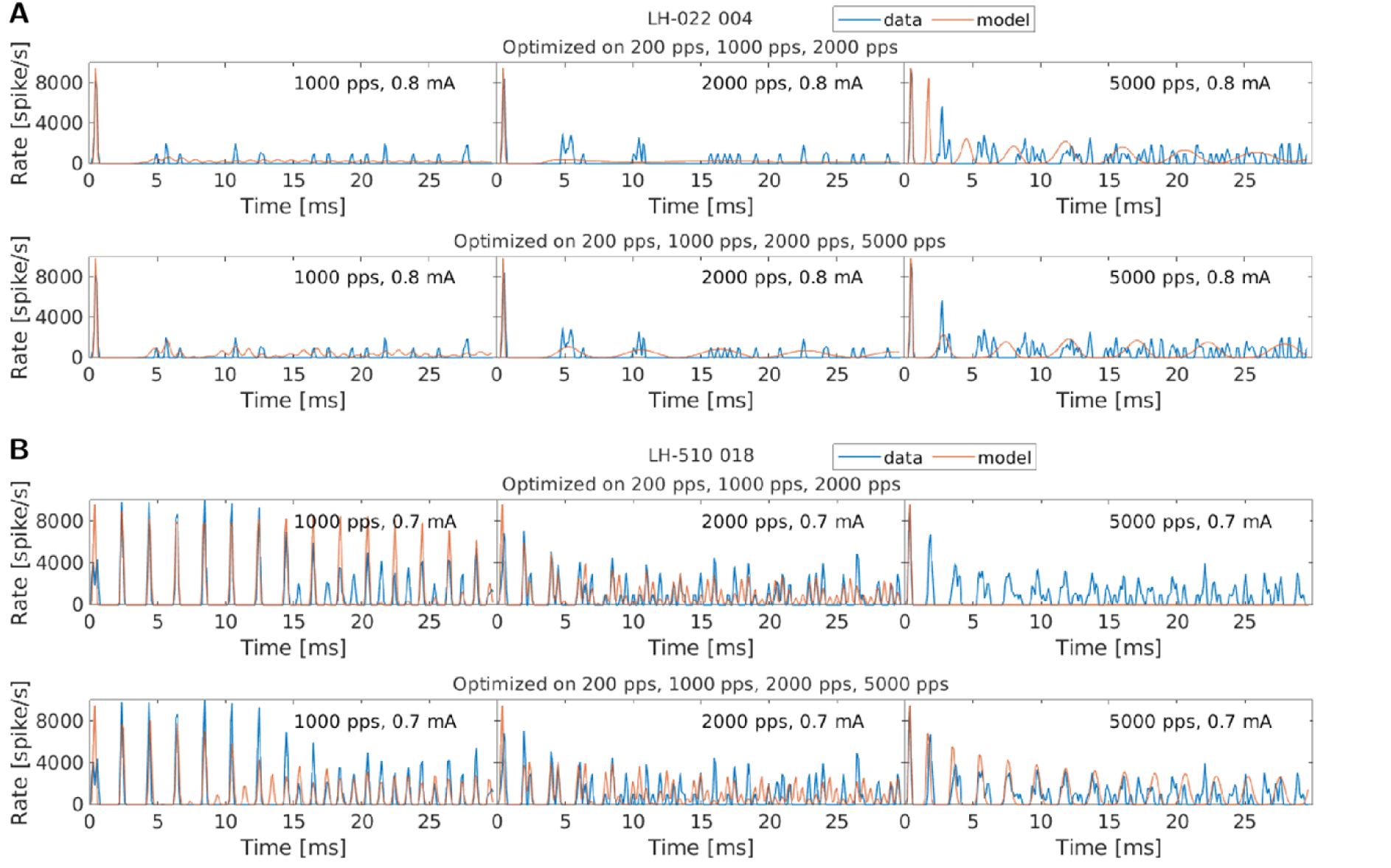
The responses of two exemplary fibers and the model responses to the three pulse rates of 1000 pps, 2000 pps, and 5000 pps at the highest stimulating amplitude. The upper row in each panel shows the results of the original optimization without the 5000 pps data. In the lower row in each panel, the 5000 pps data was included in addition to the other rates.

Figure 7 shows the three similarity terms for all 10 refitted fibers, for the original fit without the 5000 pps data in blue and for the parameter sets also fitted to the 5000 pps data in green. It shows that for the refit the similarity terms improved drastically for the 5000 pps data, as would be expected. However, Figure 7 also shows that the similarity values for the three other pulse rates became only slightly worse if the 5000 pps data is included.

**Figure 7:**
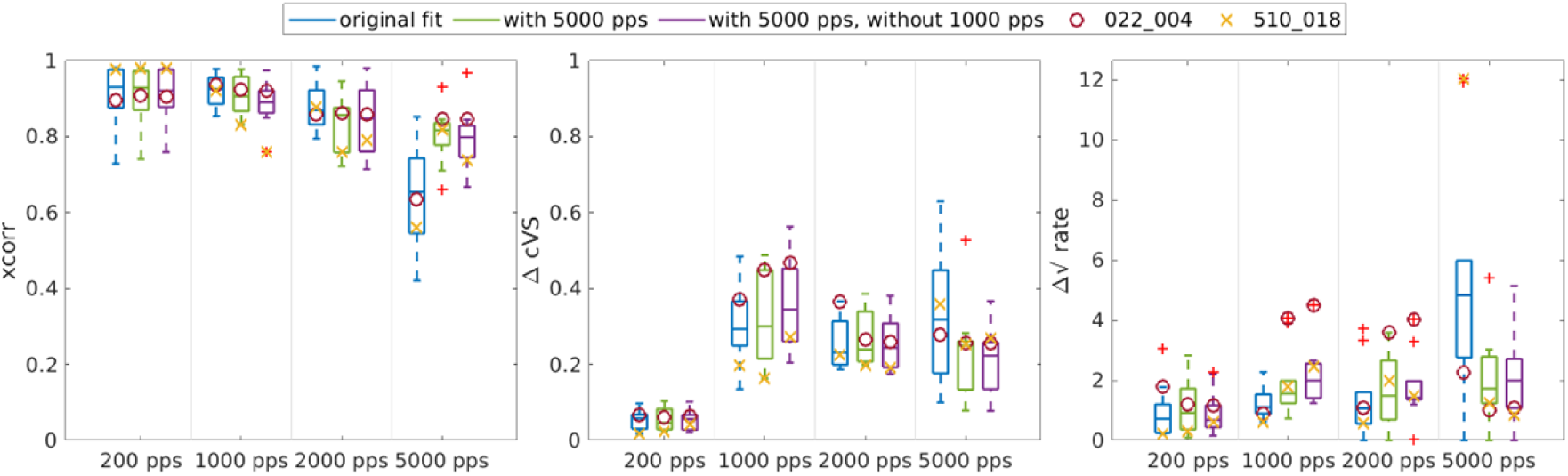
Comparison of the similarity terms for the original fit (blue), the refit with 5000 pps included (green), and the refit with the 5000 pps included but without the 1000 pps data (purple) for all 10 fibers that were reoptimized. The symbols indicate the values of the two fibers shown above.

In the previous section we have shown that there might be multiple parameter sets that led to similar simulation results. The improved simulation of the 5000 pps data, without a high decrease in performance for the other pulse rates, indicated that also for these fibers there are multiple parameter sets that led to similar results for the lower pulse rates, and including the 5000 pps data disambiguates between them. Unfortunately, we did not have the 5000 pps data available for all fibers and did therefore not include them in the main optimization to preserve the comparability of the fits to the individual fibers.

Following up on the improved fit to the 5000 pps data, the question arose if our model can only predict pulse rates that had been seen previously, or if the problem with the 5000 pps data was rather due to the fact that time points in the response functions became relevant that had not been sampled before by the lower stimulation rates. In other words, is the model limited in extrapolating, or is it even limited in correctly interpolating between pulse rates? To answer this question, we ran the optimization procedure for the same 10 fibers again, but this time included only data from the pulse rates 200 pps, 2000 pps, and 5000 pps. The similarity terms for these simulations are shown in purple in Figure 7. Comparing these similarity terms with the ones from the optimization including the 1000 pps data (in green) shows that there were only small changes in performance for all four pulse rates. Including the 1000 pps resulted in a slightly improved performance at 1000 pps but not at the other three rates. This indicated that the aLIFP model, together with the fitted parameters, can interpolate between previously seen pulse rates.

Regarding the efficiency of the optimization, it is also interesting to note that including all four pulse rates led to a much higher optimization duration of 12 ± 6 hours, and removing the 1000 pps values reduced the optimization duration again down to 9 ± 5 hours.

### Comparison of the phenomena with literature

Another step in the evaluation of our fitting was to compare the characteristic values of seven phenomena obtained from the fitted parameter sets with the values obtained from literature. The first characteristic value we computed was the dynamic range, which was calculated as the ratio in decibels between the amplitudes resulting in spike probabilities of 0.1 and 0.9 in response to a single pulse. The description of all other characteristic values can be found above in the section Parameter certainty.

In the literature, not for all phenomena values were available from guinea pigs, and even from other species (cats) only a very limited amount of data could be found, as discussed in the Introduction. An overview of the literature values used is provided in Table 1. The literature data also includes the dynamic range, latency, and jitter from the original publication of the data used in this work by Heffer et al. [9] However, as we re-detected the spike times, we also recalculated the values and provided them additionally. This changes especially the spike latency, as the original publication placed the spike at the rising slope of the action potential, while the re-detection placed the spike times at the maximum.

**Table 1:** This table gives an overview of the literature values used in Figure 8.

| Phenomenon | Source | Species | Values | Method of calculation |
| --- | --- | --- | --- | --- |
| Dynamic range | Sly et al. (2007) [28] | Guinea pig | 1.2 dB | Single pulse, provided in the paper |
|  | Miller et al. (1999) [4] | Cat | 1.38 dB | Single pulse, provided in the paper |
|  | Heffer et al. (2010) [9] | Guinea pig | 0.67 dB | First pulse of 200 pps stimulation, provided in the paper |
|  |  |  | 1.09 dB | First pulse of 200 pps stimulation, recalculated based on the data |
|  | Model based (mean) | Guinea pig | 1.01 dB | Single pulse |
| Latency | Sly et al. (2007) [28] | Guinea pig | 0.57 ms | Single pulse, provided in the paper |
|  | Miller et al. (1999) [4] | Cat | 0.61 ms | Single pulse, provided in the paper |
|  | Heffer et al. (2010) [9] | Guinea pig | 0.54 ms | First pulse of 200 pps stimulation, Provided in the paper |
|  |  |  | 0.72 ms | First pulse of 200 pps stimulation, recalculated based on the data |
|  | Model based (mean) | Guinea pig | 0.75 ms | Single pulse |
| Jitter | Miller et al. (1999) [4] | Cat | 71 μs | Single pulse, provided in the paper |
| | Heffer et al. (2010) [9] | Guinea pig | 130 μs | First pulse of 200 pps stimulation, provided in the paper |
| | | | 108 μs | First pulse of 200 pps stimulation, recalculated based on the data |
| | Model based (mean) | Guinea pig | 140 μs | Single pulse |
| Refractory period | Cartee et al. (2000) [16] | Cat | 0.77 ms | Calculated based on the paper |
|  | Miller et al. (2001) [5] | Cat | 1.08 ms | Calculated using the aLIFP model |
|  | Model based (mean) | Guinea pig | 2.36 ms |  |
| Facilitation duration | Cartee et al. (2000) [16] | Cat | 0.34 ms | Calculated based on the paper |
|  | Dynes (1996) [8] | Cat | 0.82 ms | Calculated using the aLIFP model |
|  | Model based (mean) | Guinea pig | 0.62 ms |  |
| Accommodation duration | Dynes (1996) [8] | Cat | 4.61 ms | Calculated using the aLIFP model |
|  | Model based (mean) | Guinea pig | 3.58 ms |  |
| Adaptation amplitude | Zhang et al (2007) [6] | Cat | 1.01 | Calculated using the aLIFP model |
|  | Model based (mean) | Guinea pig | 1.01 |  |

Comparing the values obtained from the model first with the values obtained directly from the data showed that the dynamic range and the latency could be very well estimated from the model. However, the jitter was on average smaller than the values obtained directly from the data. This is because a lower jitter was required to fit the vector strength accurately. This indicates that the jitter during the course of stimulation is slightly overestimated by the model.

Comparing the characteristic values of the phenomena of the nerve fibers used in this work with the literature data in Figure 8 shows that all phenomena values obtained in this work and the literature values were in a similar range. For the refractory period, the values obtained here are on average larger than the two literature values, while the accommodation is on average smaller than the 4.61 ms we obtained with our model and the data by Dynes (1996). The characteristic values of both facilitation and adaptation are very similar compared to the values calculated from literature.

**Figure 8:**
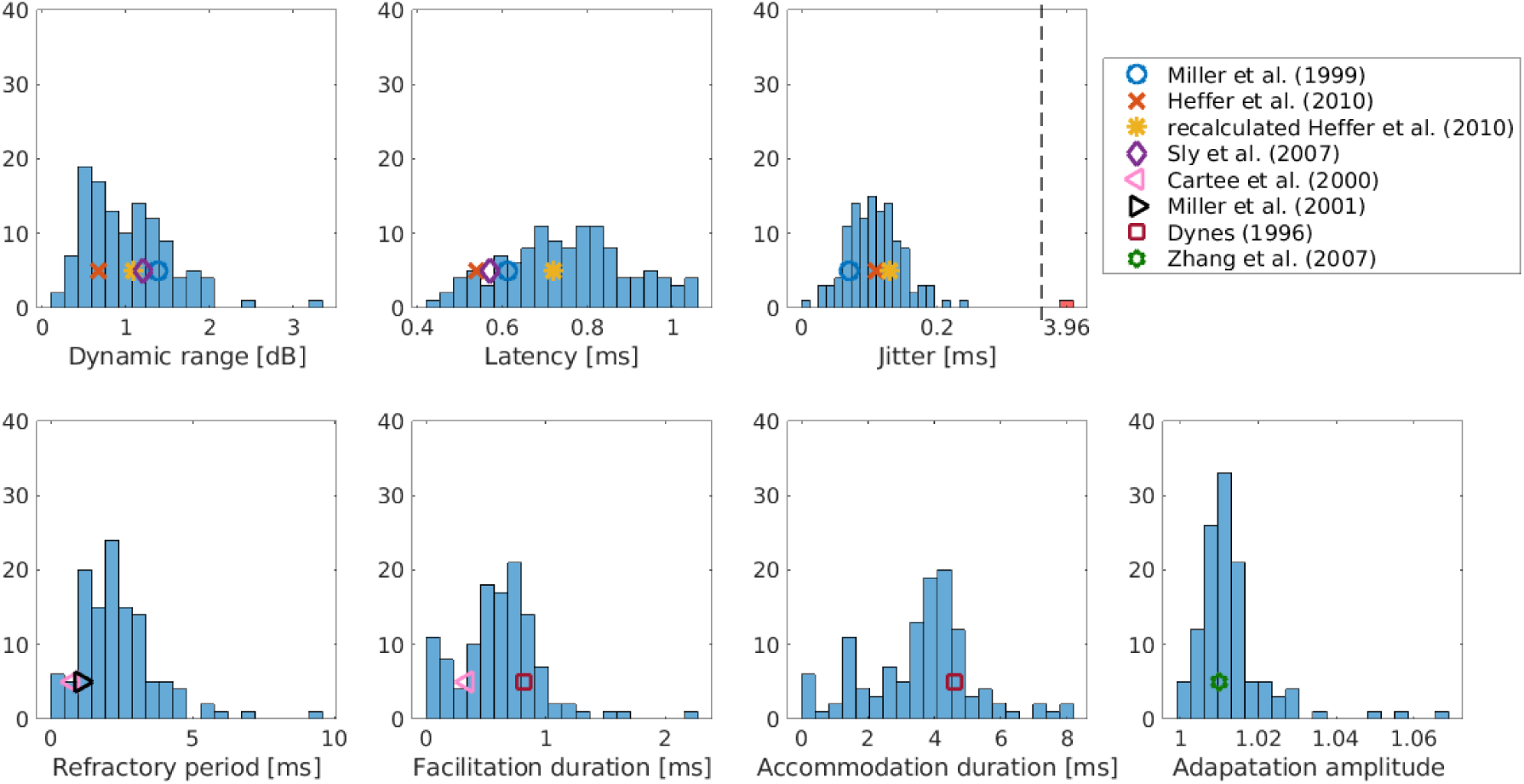
Comparison of the characteristic values of seven phenomena discussed here, with values taken or calculated from the literature.

Most literature values have been obtained with dedicated experiments that investigate one specific phenomenon. This holds especially true for the refractory period, facilitation, and accommodation. These values have all been obtained using paired pulse experiments, which investigate the change in the spike threshold after a supra-or sub-threshold stimulation. The similarity of the values obtained from pulse-train stimulation to the literature values shows that it is not necessary to perform a dedicated experiment for each phenomenon. Instead, the analysis of pulse-train responses with a detailed model, such as the aLIFP model, is sufficient to provide realistic estimates for the phenomena.

### Phenomena interaction

After evaluating the reliability and realism of our fits, we wanted to demonstrate a possible application of our work. For this, we went back to the example of Marder and Taylor [17], which we already discussed in the Introduction. They argued that possible interactions between different measured values get lost if only population mean values are compared. Here, we obtained the characteristic values for seven phenomena for each nerve fiber and computed the cross-correlation between all characteristic values (Figure 9). For most pairs of phenomena, the correlation was relatively low, however, we want to discuss the five pairs with an absolute correlation larger than 0.3.

**Figure 9:**
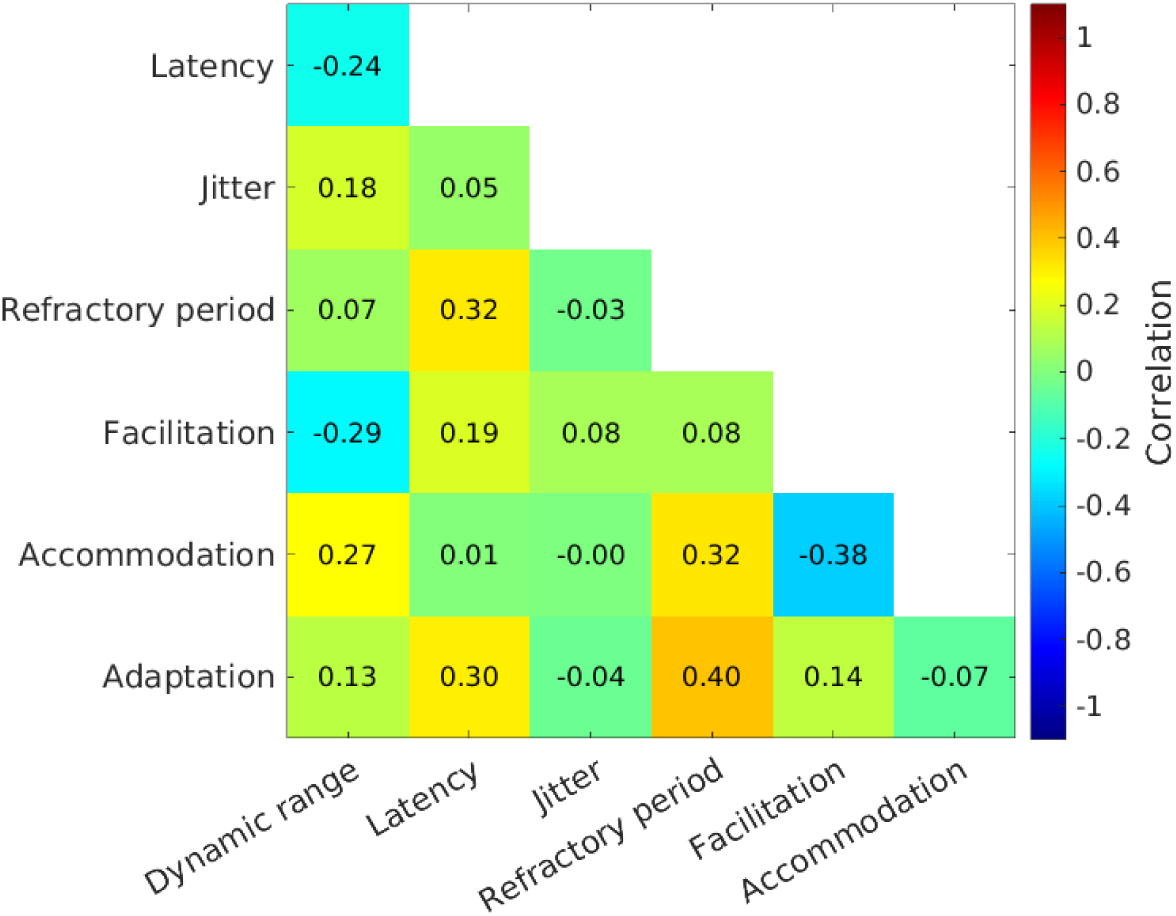
The correlation between the characteristic values of all seven phenomena considered in this work. In addition to the color coding, also the exact values of the respective pair are shown.

**Figure 9:**
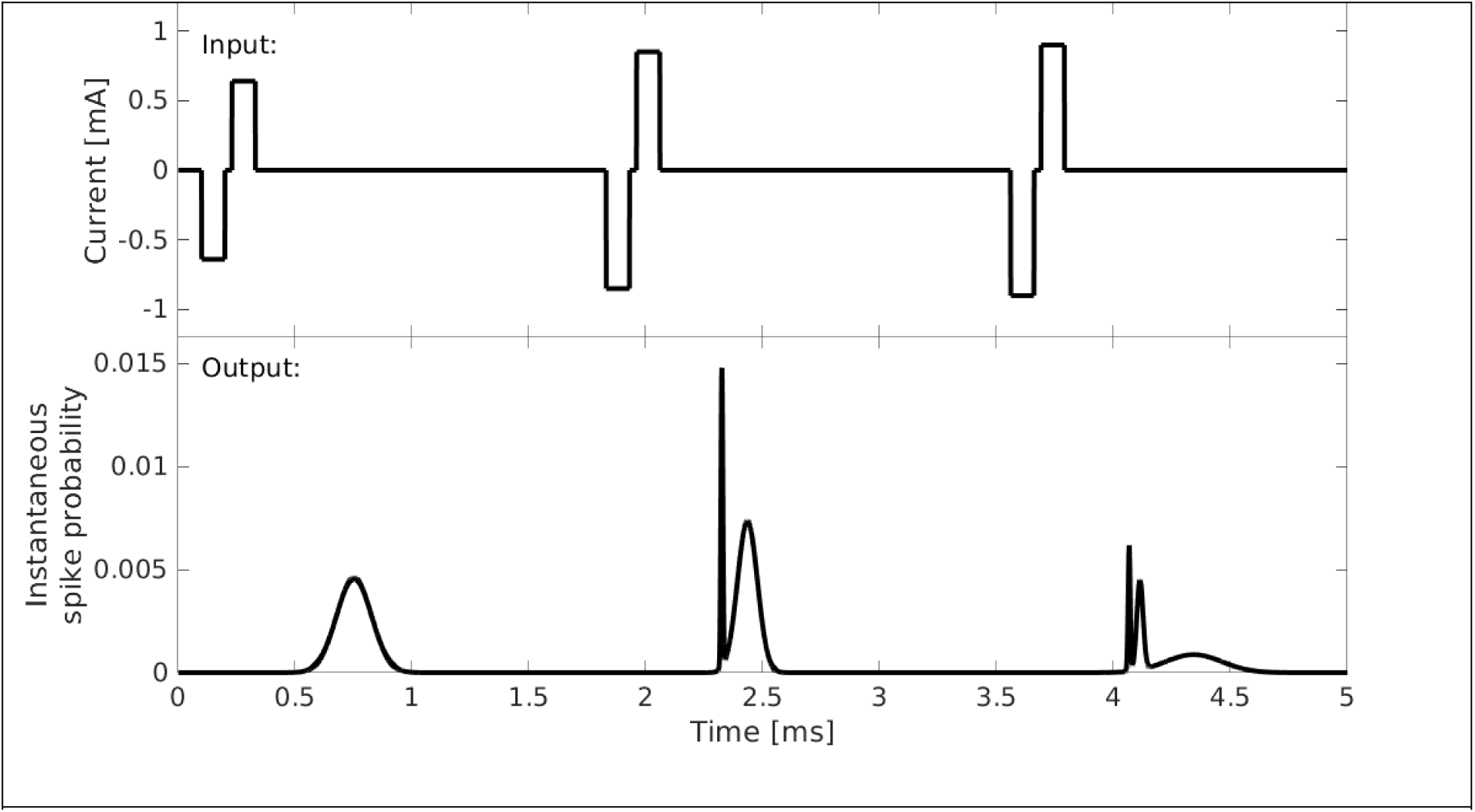
An exemplary output of the aLIFP model. The upper panel shows the input, which consists of three bi-phasic pulses with different amplitudes. The lower panel shows the output of the model, which is the instantaneous spike probability over time.

The refractory period was positively correlated with three other measures: the latency at 50% spike probability, the accommodation duration, and adaptation amplitude. All four parameters could be related to the membrane time constant, or the speed of the ion channels, and a correlation is therefore biophysically plausible. Also, Negm and Bruce [12] theorized in a similar direction by arguing that the same set of ion channels might cause all pulse interaction phenomena. A dedicated experiment to investigate this correlation might be very interesting. In the light of this theory, it is interesting to note that, except for latency and adaptation, the four phenomena correlated only with the refractory period, not with each other. The most probable reason for this is the uncertainty in the estimated parameters described above. As the correlation was only small itself, even small errors in the parameter estimation can counteract the correlation completely.

The last correlation we highlight was a negative correlation between the accommodation duration and facilitation duration. With the argument of a membrane time constant, which affects all timing-related phenomena, a positive correlation between these two phenomena would be expected. However, both sub-threshold phenomena were described using a single function in the aLIFP model. Figure 10D illustrates that the two fitted parameters can affect both facilitation and accommodation, and longer accommodation could easily lead to shorter facilitation and vice versa. We therefore cannot assume that the facilitation and accommodation duration in the aLIFP model are independent, and the negative correlation found here was very likely only due to the underlying model.

**Figure 10:**
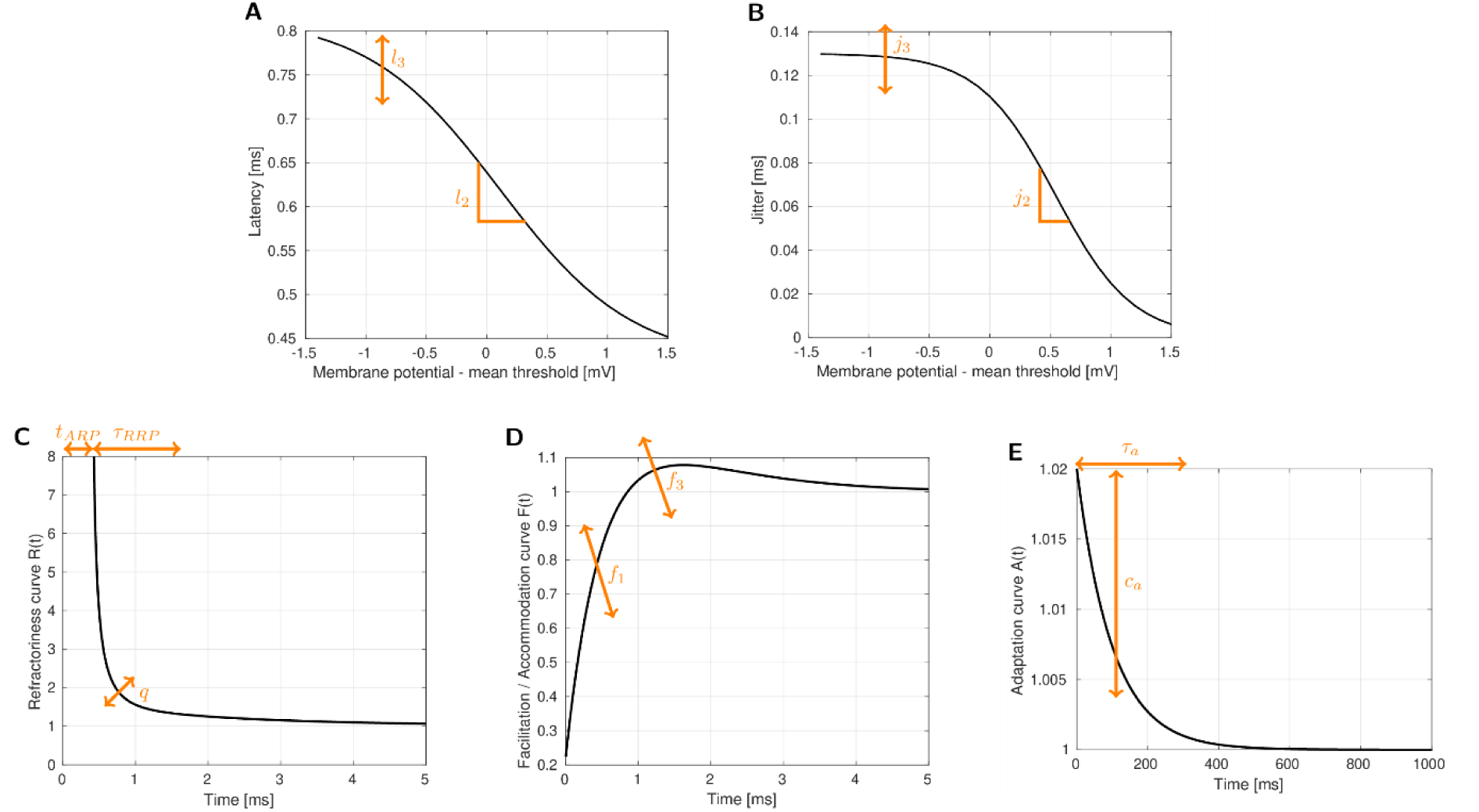
An illustration of how the optimized parameters affect the curves describing latency and jitter (A and B) and the pulse interaction phenomena (C–E). Note also the longer time scale of the adaptation curve compared to the refractory curve and facilitation/accommodation curve.

## Discussion

In this work, we showed that it is possible to optimize the parameters of the aLIFP model such that the behavior of individual nerve fibers is simulated in detail. Thus many different response patterns were accurately replicated. Based on the fitted parameter sets for each nerve fiber, we extracted characteristic values for seven phenomena. We show that these values were in the same range as values obtained from the literature, which further validated our method.

### Optimization procedure

The success of every optimization procedure depends very much on the chosen cost function, which describes the similarity between the target (data) and the output (model simulations). Here, choosing the cost function was particularly difficult, because it had to satisfy two contradictory conditions. On the one hand, the cost function should describe the difference between the data and the model as detailed as possible, ideally such that the procedure is able to fit the data with a temporal accuracy well below 1 ms. On the other hand, the cost function had to be robust against the stochasticity in the fiber response. This stochasticity was especially detrimental at low response rates, even though the deterministic description of the stochasticity in the aLIFP model greatly facilitated the fitting. We argue that such a detailed fit was only made possible by this deterministic description. Nevertheless, combining the two aims with the cost function was not a straightforward task. The solution was a cost function that consisted of three terms that differently expressed the similarity between the data and the model simulations: cross-correlation, complex vector strength, and rate difference. The cross-correlation of the first half of the responses allowed detailed temporal fitting that was supported by the complex vector strength difference. In the second half of the response, we calculated the difference between the square roots of the rates. This was not sensitive to detailed differences in the response pattern but robust against the noise that was more prominent in the second half of the response.

However, this cost function is non-convex, which made it very difficult to find the global minimum. We therefore chose the solver BADS for our optimization problem, which is able to deal with such a cost function [29]. BADS came with the additional benefit that it could speed up the optimization of computationally expensive cost functions by predicting intermediate responses based on Bayesian optimization. However, as the aLIFP model still had to be run thousands of times, we split our optimization problem into several steps to perform the optimization in a reasonable time. Even though BADS can deal with a complex optimization landscape, we still had to start the third optimization step from several starting points to reliably converge to a good solution. The starting points were manually selected to result in a wide variety of response patterns. However, optimizing parameters sequentially can lead to a globally sub-optimal solution, and an optimization procedure that is able to optimize all parameters at the same time would have been preferable.

### Generalization and model design

In an ideal case, the parameters fitted to a small amount of recorded data of a nerve fiber have general validity, i.e. they can predict response patterns to other stimuli. We showed that it is possible to predict unseen response patterns at intermediate pulse rates, but the model responses do not extrapolate to unseen higher pulse rates (Figure 7). From the point of view of the aLIFP model, this can be explained by an under sampling of the functions describing the phenomena. This under sampling was also the most probable cause of the parameter ambiguity that we described in Figures 4 and 5. The underlying problem for both the lack of extrapolation and the parameter ambiguity is the very phenomenological approach of the aLIFP model, where each phenomenon is described separately by a function. This is the case even though some phenomena, such as latency, refractory period, or accommodation, are most likely all caused by the same biophysical mechanisms: the ion channels or membrane characteristics. It is therefore biophysically implausible to model them as independent phenomena in a model. The separate description of the phenomena was also induced by the existing literature, which investigated most phenomena separately, resulting in the impression of independence. Even though some phenomena are regularly set in relation to each other (e.g., latency and spike probability in Miller et al. [4]) and are thus also thus modeled, the relationships between the different response phenomena have not been systematically investigated. In Figure 9, we showed that the aLIFP model can capture this covariance of some of the phenomena if the parameters are fitted to enough data. However, due to the many degrees of freedom in the model, the parameter sets found were ambiguous. A model that constrains the parameters in a biophysically plausible way could therefore facilitate the fitting of individual nerve responses. This could, for example, be a membrane time constant that links latency, refractory period, facilitation, accommodation, and adaptation. However, it is also important to note that to find such biophysically plausible constraints, an understanding of the interaction between the response phenomena is necessary, as it can be obtained by fitting the behavior of individual nerve fibers. Based on such knowledge, models can be updated or a new model created.

### Model limitations

The optimization procedure developed provided us with the possibility to test the abilities and constraints of the aLIFP model in a very detailed manner. We found that the model can fit many different response patterns with remarkable accuracy, as shown in Figures 1 and 2. However, it is also interesting to look at the patterns where the model struggles to simulate them accurately.

Of the 20 fibers for which the model simulation had, on average, the lowest correlation with the data, 14 fibers had a very similar pattern. In these fibers, a high, modulated response after the first pulse at medium-to-low stimulation amplitudes was seen (e.g., Figure 1b fiber LH-011_012), which cannot be simulated by the model. We tried to adjust the model parameters manually to model this, but did not succeed, which indicated that this is a limitation of the model, and not of our optimization procedure. A more detailed analysis of the problem showed that following the first pulse, a larger threshold standard deviation, or higher dynamic range, would be necessary to reproduce this behavior.

Another phenomenon that was only poorly simulated by the model is the transition from a modulated response to a more stochastic, unmodulated response. Both behaviors could be replicated if they occur separately, but the optimization procedure had problems fitting parameters that could replicate this transition. Adjusting the model parameters manually showed that although the model can replicate the transition accurately, this does not emerge naturally from the parameters fitted to the first half of the responses. Thus our cost function, which only considered the rate difference in the second part of the response, was not ideally chosen. A cost function that also considers those details of the response could lead to parameters accurately describing this transition. However, such a cost function also needs to be able to deal with the increased stochasticity in the second half of the response.

### Fit duration

Fitting the aLIFP model with the proposed optimization procedure was computationally expensive. This was the case even though the response can be obtained from a single run, and we took multiple steps in designing the optimization procedure to reduce the computational cost. The original fit still took on average 8 ± 4 hours per fiber to optimize the parameters on a single core of a Xeon Gold 6238 CPU. If the 5000 pps were also included, it took 12 ± 6 hours. Due to the long optimization duration, it is therefore important to carefully balance the benefits of more stimulus conditions, which improve the quality of the fit, against the fitting duration. As we showed above, not all data led to a similar improvement in fitness. Including the 5000 pps condition improved the simulation of this condition drastically, but removing the 1000 pps condition only led to a very small decrease of the similarity metrics. It is therefore an important part of the fitting procedure to carefully design the dataset to which the parameters are fit. The proposed optimization procedure can also be used to design informative experiments, as was also proposed by Pozzorini et al. [24], who used models to suggest or even select the most informative measurement points.

### Observations during the optimization

Solving and understanding the problems that occurred in describing certain nerve fibers with the aLIFP model also forced us to look very closely at the dataset. This then led to discoveries in the data that might have gone unnoticed without the model-based analysis, even though every experimenter tries to understand the data as well as possible. The first discovery we made is a slight deviation from the intended pulse rate for four nerve fibers. This was only discovered due to a very slight beating or latency drift between the simulated pulse rate and the recorded one. For most applications, such a small deviation is not relevant, but it can make a difference when, e.g., analyzing the data with a model or computing the vector strength.

As we looked closely at the fibers for which the model could not accurately simulate responses, we found several fibers that became less responsive over the course of multiple measurement repetitions. This is a behavior that the model cannot predict, as the response characteristics do not change during the simulation.

Similarly, we also detected more cases where the responses of two nerve fibers were recorded than we had detected manually. This happened if there were two responses to a single pulse, e.g., at 200 pps.

### Further applications

There are several possible applications for this optimization procedure. As we have illustrated, it can be used for the analysis of datasets on a fiber-by-fiber basis. In such an analysis, it can provide more information than a manual analysis, as the optimization is able to estimate all phenomena shaping the fiber response. In this way, hypotheses about individual nerve fiber behavior or the correlation between phenomena can be generated, which can then be tested in dedicated experiments. Such an analysis can also be taken well beyond the present work, e.g., by comparing different sub-populations in the datasets, or by identifying groups of fibers that share similar parameters.

In a similar way, as noted above, the procedure can be used to decide on the most informative stimuli for new experiments. This can be carried out by testing the responses to which stimuli can be predicted from the training data, and which responses cannot. In our case the 5000 pps stimuli were much more informative than the 1000 pps stimuli.

The procedure can also be used to test existing models, as here with the aLIFP model. In this way, one can see if different nerve fiber behaviors can be predicted by a single model just by changing its parameters. Problems in simulating certain behavior can then lead to questions about the knowledge on which the model is based, or to questioning the model itself. Thus both the models and the experimental paradigms can be improved, as discussed above.

Not only can models be tested, but also the quality of datasets. To be able to predict a dataset with a model, the data needs to be carefully recorded and labeled, and any inaccuracies will become apparent when trying to simulate the behavior of the nerve fibers on an individual basis.

## Conclusion

We proposed an optimization procedure that finds 118 nerve-fiber-specific model parameter sets based solely on pulse-train stimulation, such that a wide range of response patterns can be simulated. The resulting parameter sets can then form the basis for a detailed analysis of data on a fiber-by-fiber basis. To illustrate this, we showed correlations of the duration of the refractory period with latency, accommodation duration, and adaptation amplitude. However, when drawing conclusions from such a model-based analysis, it is important to be aware of the underlying model structure and its limitations. The procedure proposed can provide valuable insights, and can aid in defining hypotheses and stimuli for further experiments. Furthermore, it can serve as a basis for the development of model-based fitting of nerve fiber responses and, ultimately, contribute to a more detailed understanding of the electrode-to-nerve interface in neuroprosthetics.

## Methods

### Data

The data used in this work was originally collected as part of the work of Heffer et al. [9]. They recorded in-vivo responses from 188 electrically stimulated auditory nerve fibers of 25 adult guinea pigs. 15 guinea pigs were acutely deafened, and 10 guinea pigs were deafened five weeks prior to the experiment. In this work we do not distinguish between acutely deafened and five-week deaf fibers. For details of the deafening and recording procedure, please refer to the original publication [9].

The stimuli were 100 ms constant-amplitude pulse trains with bi-phasic pulses, which had a phase duration of 25 µs and an inter-phase gap of 8 µs. The pulse rates were 200 pps, 1000 pps, and 2000 pps, and for 97 fibers the response to 5000 pps pulse trains was also recorded. All fibers were stimulated with multiple amplitudes across their dynamic range, with an average of 11 ± 5 amplitudes and 24 ± 16 repetitions per condition.

In [9], the spike detection was based on a thresholding procedure, which is prone to errors [30]. We have therefore re-detected the spike times using template matching. A description of the spike detection procedure can be found together with the voltage traces and detected spike times at Zenodo (DOI: 10.5281/zenodo.15827116).

From the 188 nerve fibers that were used in the work of [9], we have omitted 39 due to problems during spike extraction (13 due to a noisy recording and 26 due to stimulation of two fibers). During the optimization of the parameters, we removed a further 15 fibers, as either not enough repetitions or not enough stimulation amplitudes with responses were available. Another 10 fibers were removed due to a systematic reduction in the response over the course of multiple repetitions of the same stimulus, and 6 fibers were removed due to probable double spiking, which had not been detected during the spike detection procedure. This left us with 118 fibers for the optimization.

While trying to describe the data with the aLIFP model, we noticed for some fibers a slight beating or latency drift between the model response and the data. The only way to explain this was a slight inaccuracy in the stimulating pulse rates. We therefore detected the stimulating rate from the artifact in the recording. From this detection we found a deviation of the intended pulse rate for four fibers, with a maximum deviation of the pulse rate of 9%. As deviations were partially detrimental for the fitting success, we always used the estimated pulse rate for the model stimulation. However, to simplify the description, we will continue to use the intended pulse rate when referring to a pulse rate.

### Neural model

The model fitted to the neural data was the adaptive Leaky-Integrate and Firing Probability model (aLIFP; [27]). Instead of predicting individual stochastic spike times, the model predicts the spike probability over time. In this way, the stochastic nature of the spike times can be described by a fully deterministic model. For the task of fitting stochastic spike times, this provided the advantage that only the stochasticity of the data must be considered. An exemplary model output can be seen in Figure 9.

As the name indicates, the aLIFP model is based on a leaky integrator, which transforms the input current into a passive membrane potential. In a leaky-integrate and fire model a spike is evoked once a threshold is crossed [14]. To describe the noisiness of the fiber response, the threshold is often modeled by drawing from a Gaussian distribution in each time step. In the aLIFP model, instead of drawing from the distribution, all calculations are directly performed on this distribution. Through this, a distribution for the time point of threshold crossing is obtained, which is also described using a Gaussian. The resulting distribution of the expected spike times is then a Gaussian distribution as well. It is calculated by combining the distribution of the time point of threshold crossing with the latency and jitter. Latency is the mean duration between the stimulation time and the spike time, while jitter is the standard deviation of the latency. Both latency and jitter are dependent on the total spike probability and are larger for a low spike probability and smaller for a high spike probability [27]. In the aLIFP model, the latency function is controlled by four model parameters, from which two were optimized, and by the three-parameter jitter function, for which another two parameters were optimized. The parameters controlling the shift on the x-axis (l_1_ and j_1_) have been set to 0, as only a limited amount of data was available below a spike probability of 0.5. Setting the value to zero forces the function to be symmetrical around a spike probability of 0.5. For the additional offset parameter l_4_ of the latency the default model value was used [27]. For both latency and jitter, one of the optimized parameters controls the slope of the function (l_2_ and j_2_), while the other controls the maximum amplitude (l_3_ and j_3_). The effect of the two parameters is illustrated in Figure 10 in panels A and B.

The aLIFP model includes the three short-term pulse interaction phenomena: refractory period, facilitation, and accommodation and the long-term pulse interaction phenomenon adaptation. All four phenomena are described by functions, which are multiplied with the model threshold. The refractory period is the decrease in excitability after a spike was evoked. It can be divided into the absolute refractory period, where spiking is not possible, and the relative refractory period, where a higher stimulus amplitude is needed to cause a spike. In the aLIFP model, the refractory period is described by an increase in the phenomenological threshold. The duration of the absolute refractory period is defined by t_ARP,_ and the duration and shape of the increase during the relative refractory period are parameterized by τ_RRP_, p, and q [27]. To reduce the number of parameters used for fitting, only t_ARP,_ τ_RRP,_ and q were changed here. In Figure 10C the shape of the refractory function is shown, together with the influence of the three parameters.

Facilitation and accommodation occur if a pulse was not able to evoke a spike. During facilitation, the excitability of the nerve fiber is increased, while the following accommodation decreases the excitability. In the aLIFP model, this is again caused by changes in the model threshold. The changes to the threshold are described by a function with five parameters [27]. However, the influence of the parameters on the curve interacts, and to facilitate the fitting, only two of the five parameters were fitted. The parameter f_1_ mainly controls the amount of facilitation, while f_3_ controls the amount of accommodation, which is illustrated in Figure 10D.

Adaptation is the long-term decrease in excitability after multiple spikes occurred, which cannot be explained by refractoriness alone. In the aLIFP model, it is described by a slight increase in threshold after every spike. The duration of the increase is controlled by the time parameter τ_a_, and the amplitude of the increase by c_a_. This increase is illustrated in Figure 10E, together with the influence of the two parameters. Note the larger time scale in comparison to the functions describing the refractory period and the facilitation and accommodation. The third adaptation parameter of the aLIFP model, which describes the maximum adaptation amplitude possible, was not fitted due to its low influence on the model output during a time scale of less than one second.

In total, the model has 23 parameters of which 13 were fitted in this paper. As the data used in this work was obtained using a single pulse shape and pulse duration, the time constant of the leaky integrator and the duration of the action potential initiation period could not be found. Also, the number of parameters fitted for the interaction phenomena has been reduced, as explained above.

To aid the fitting and to achieve a meaningful result, we limited the value range of most of the parameters. Table 2 gives an overview of all fitted parameters of the aLIFP model together with their respective ranges. As the parameters for the facilitation curve strongly interact with each other, placing limits on them was not enough to achieve a meaningful result. We therefore placed an additional constraint on the facilitation curve, that the first value must always be below 1.

**Table 2.**
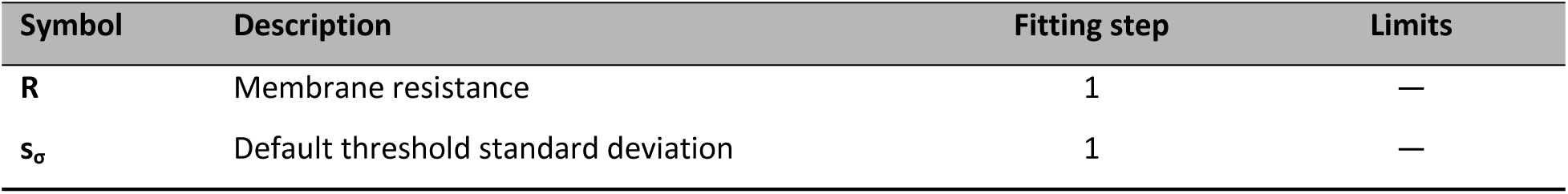
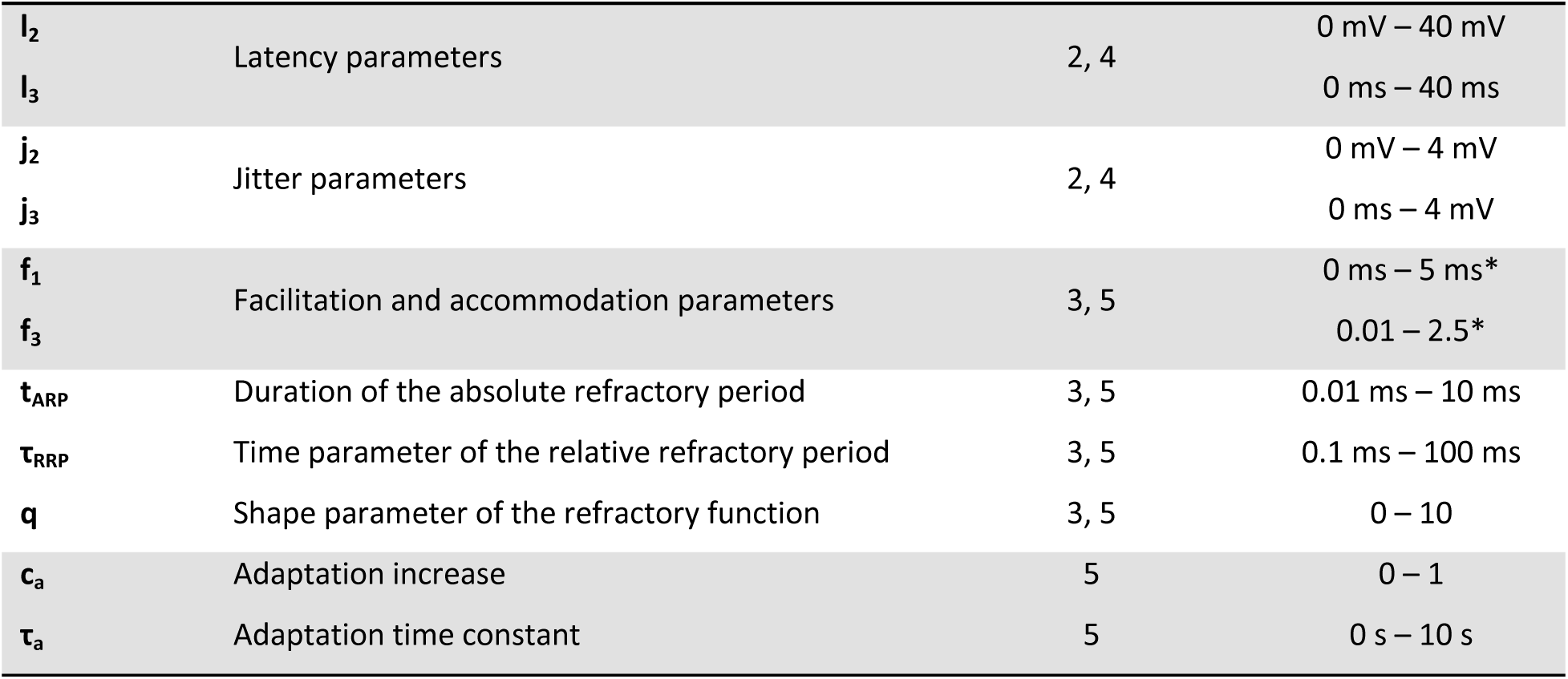
This table gives an overview of the parameters of the aLIFP model used during the optimization and states the optimization steps where they were optimized together with their limits used during the optimization. On the facilitation parameters (marked with *), an additional constraint was placed, such that the first value of the facilitation function is always below 1.

### Fitting procedure

The aim of the fitting procedure is to find parameters of the aLIFP model such that the model output describes the spiking pattern of a single nerve fiber as well as possible. Due to the many phenomenological details of the spiking process included in the aLIFP model, fitting the parameters was not a straightforward task, and a mathematically closed solution was not possible. Therefore, a numerical fitting procedure consisting of multiple steps was developed. In the following, first the cost function used in most of the fitting steps will be presented, and then the five individual steps. An overview of the five steps of parameter fitting can be found in Table 3.

**Table 3.** Summary of the individual steps of the optimization procedure.

| | Description | Pulse rates | Duration | Cost function | $C_{xcorr}$ | $C_{cVS}$ | $C_r$ |
| --- | --- | --- | --- | --- | --- | --- | --- |
| 1 | Fitting of membrane resistance and threshold standard deviation | 200 pps | First pulse | Least-squares | – | – | – |
| 2 | Fitting of latency and jitter parameters | 200 pps | First pulse | Least-squares | – | – | – |
| 3 | First fit of short-term interaction parameters | 200 pps, 1000 pps, 2000 pps | First 20 ms | Eq. (8) | 1 | 0 | 0 |
| 4 | Refit of latency and jitter parameters | 200 pps, 1000 pps, 2000 pps | First 20 ms | Eq. (8) | 1 | 0.5 | 0 |
| 5 | Refit of short-term interaction values, fit of adaptation parameters | 200 pps, 1000 pps, 2000 pps | All 100 ms | Eq. (8) | 3 | 0.75 | 0.1 |

#### Cost function

The cost function of fitting steps 3, 4, and 5 consisted of three similarity terms that describe the difference between the model and the data response. The first term was the normalized cross-correlation between the response of the data r_d_ and of the model r_m_. The aim of this part was to describe the similarity of the envelope of the response of the model and the data. Due to the stochasticity of the biological spike generation, calculating the cross-correlation provided only meaningful results if the number of spikes did not become too low. As the number of spikes decreases with increasing signal duration, we calculated the cross-correlation only in the first part of the signal. The duration of the first part of the signal was obtained for each condition of each fiber separately, as the speed of the response decrease varied with condition and fiber. The duration of the first part of the signal was defined to be at least 20 ms and at most 50 ms. During this interval, the number of spikes across all repetitions of one condition was counted in non-overlapping windows of 10 ms. Once the number of spikes per window was below 10, the individual end point was found. The earliest end point was 20 ms, and if the number of spikes did not decrease below 10, 50 ms was used as the end point of the first part of the signal.

The basis for the calculation of the cross-correlation was the time dependent spike rate. This rate was obtained in windows with a total length d_w_=0.6 ms. The windows had linearly rising and decaying slopes of 0.2 ms and a constant plateau of 0.2 ms. Two consecutive windows overlapped by 2/3 d_w_. The instantaneous model rate in the i*th* window was obtained from the spike probability P_s_(t) as

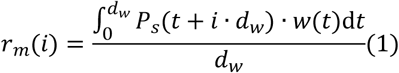

The instantaneous data rate r_d_ in the i*th* window was calculated from the time point s_n_ of the spikes as

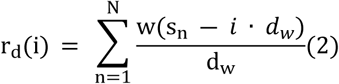

where N is the total number of spikes. The first part of the cost function was then obtained as the normalized cross correlation between the r_d_ and r_m_:

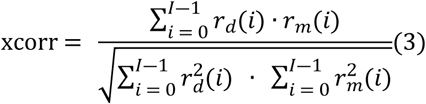

where I is the total number of windows.

The aim of the second similarity term in the cost function was to describe both the similarity of the phase locking and the similarity of the response phase between the model and the data. For this we used a variation of the vector strength, which we call the complex vector strength. The vector strength was originally introduced by Goldberg et al. [31]. They represented the phase relation between each spike and the stimulating signal using the complex number

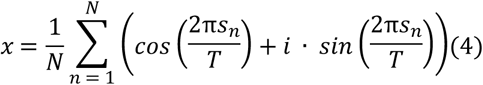

where T is the phase duration of the stimulating signal and N the total number of spikes in the responses to all repetitions of the same stimulus. The length of this resulting vector in the complex plane is called the vector strength (VS) and describes the strength of the phase locking

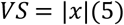

However, the argument of x contains additional information about the phase position of the spikes relative to the stimulating signal, which is discarded in the vector strength. The combination of both the length and the angle of the complex vector is here called the complex vector strength (cVS). The complex vector strength has been inspired by the complex correlation coefficient of Enke and Dietz [32].

We obtained the complex vector strength for the whole response to each condition, both for the model and the data. The second part of the cost function is then the absolute value of the difference between the complex vector strength of the model x_m_ and of the data x_d_

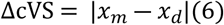

The third and final similarity term in the cost function was the rate difference between the model and the data in the second half of the response. The aim of this value was to describe the goodness of fit in the steady state of the stimulation. As this behavior was in most nerve fibers dominated by stochastic firing, calculating the correlation between the data rate and the model rate would not be meaningful here. Therefore, we used the difference of the total rate in the second half of the model, which started at the time point where the first part ends. The rate difference was then obtained as

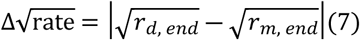

Where r_d, end_ was the total rate of the second half of the data response, and r_m, end_ the rate in the second half of the model simulation. We compare the square roots of the rates as a compromise between absolute difference and relative difference. Under the Poisson assumption that rate variance is equal to the mean rate, which is not always true in our data, a rate difference equal to the square root of the rate is also proportional to an ideal observer’s ability to discriminate the two rates [33], making it a meaningful cost function here.

Finally, all three terms were combined to produce the total cost

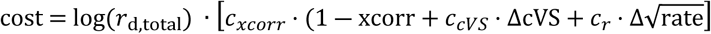

Each part of the cost function was weighed by a factor that changed for the different optimization steps. Note the subtraction from 1 for the cross-correlation, which was necessary, as the cross-correlation was maximized while the other terms were minimized. Before averaging over all conditions, the cost was weighted with the logarithm of the total rate in the data. This was done, as the conditions with a higher rate are less influenced by noise and therefore more reliable. An overview of the weighting factors can be seen in Table 3.

#### Step 1: rate-level function

In the first step, the rate-level function, which describes the relationship between the stimulation current and the spike probability (or spike rate) of the first pulse, was fitted. For this, the spike probability in response to the first pulse for each stimulation amplitude was extracted from the responses to the first pulse of the 200 pps data of the current fiber. Then the membrane resistance R and the default threshold standard deviation s_σ_ of the aLIFP model were adjusted to fit the data using a least-squares cost function. The minimization was performed using the lsqcurvefit function of MATLAB. For some fibers the dynamic range was below the difference between two stimulation amplitudes. In these cases, we increased the threshold standard deviation after the optimization to achieve the largest possible dynamic range without changing the simulation of the rate-level function.

#### Step 2: latency and jitter

Not only is the spike rate reduced when a nerve fiber is stimulated with a reduced amplitude, but also the latency between the stimulation and the spike and its jitter are affected. In the aLIFP model, this behavior is controlled by a total of seven parameters, from which four were fitted here. For this, the latency and jitter were also computed from the response to the first pulse of the 200 pps data. Then the model parameters were also found by minimizing a least-squares cost function with the lsqcurvefit function of MATLAB.

#### Step 3: short-term pulse interaction

After the parameters defining the response to the first pulse were found, we addressed the short-term interaction between individual pulses. This short-term interaction consists of the refractory period, facilitation, and accommodation. They were described in detail in the section about the aLIFP model. Only five of the nine parameters describing the short-term interaction were fitted to reduce the dimensionality of the problem. The parameters were fitted to the first 20 ms of all amplitudes and the three pulse rates (200 pps, 1000 pps, and 2000 pps) using the cost function as given in Eq. (8) with c_xcorr_=1, c_cVS_=0, and c_r_=0. Thus, this step optimized only the cross-correlation. As the optimization was highly non-convex and the computational cost of obtaining the model output for each parameter set anew was high, we used Bayesian adaptive search (BADS, [29]) to optimize the parameters. The success of the BADS can be dependent on the starting point, and we have therefore manually pre-selected six starting points for the optimization, which led to different patterns in the responses. The BADS optimization was then performed for all six starting points, and the one with the lowest cost was used for further optimization.

#### Step 4: latency and jitter

During the development of the optimization procedure, we noticed that the estimate of the jitter found in the second optimization step was often not ideal, probably due to the limited number of repetitions available. Therefore, we included another step in the optimization to refine the parameters describing the latency and the jitter. To be able to fit these to longer segments of the recordings, the optimization was done after finding first parameter estimates for the short-term interaction values.

The refit of the latency and jitter was done on the same data as used in the third optimization step, which were the first 20 ms of the responses to 200 pps, 1000 pps, and 2000 pps, and all stimulation amplitudes. For this optimization step, the cost function in Eq. (8) was used again, this time with c_xcorr_=1, c_cVS_=0.5, and c_r_=0. The cost function was chosen to also contain the cross-correlation as a regularizing term, because a perfect fit of the complex vector strength difference could also be found for completely different response patterns. Again, BADS was used as the optimization algorithm, with the previously obtained values for latency and jitter as starting points.

#### Step 5: short– and long-term pulse interaction

In the last optimization step, the five parameters controlling the short-term pulse interaction phenomena were optimized again, in addition to the two parameters controlling the adaptation. This time the optimization was performed on the complete 100 ms of recordings available, on 200 pps, 1000 pps, and 2000 pps, but only on four amplitudes, spread across the dynamic range of the neuron. The reduction of amplitudes used in this step was necessary to limit the computational time. As the optimization function, again Eq. (8) was used, with c_xcorr_=3, c_cVS_=0.75, and c_r_=0.1. The lower weight for the rate difference was necessary, as its values are not limited and can easily reach values up to 10, while the values of the cross-correlation and the complex vector strength difference are limited to a range from 0 to 1. Again, BADS was used as the optimization algorithm. The starting point for this last optimization step was the previously obtained values for the short-term pulse interaction parameters and the default values of the model for the adaptation parameters.

## Data availability statement

The nerve fiber data used in this work was published on Zenodo (DOI: 10.5281/zenodo.15827116). All code written for this paper is also available on Zenodo (DOI: 10.5281/zenodo.15848231)

## Author contributions

Rebecca C. Felsheim: Conceptualization, Data Curation, Formal Analysis, Investigation, Methodology, Software, Validation, Visualization, Writing (Original Draft), Writing (Review and Editing)

Mathias Dietz: Conceptualization, Funding Acquisition, Investigation, Methodology, Validation, Resources, Writing (Review and Editing)

David J. Sly & Stephen J. O’Leary: Data Curation, Investigation, Writing (Review and Editing)

## Notes

### Competing Interest Statement

The authors have declared no competing interest.

